# Host-derived lipids license proliferation of sterol-auxotrophic tick cells and promote flavivirus output

**DOI:** 10.64898/2026.05.11.724467

**Authors:** Helena Frantová, Alexey Bondar, Luise Robbertse, Maria Szomek, Sára Kropáčková, Arian Ebrahimi, Lucie Řimnáčová, Emanuel Barth, David Hartmann, Monika Čížková, Jan Kamiš, Lesley Bell-Sakyi, Petr Kopáček, Zdeněk Franta, Veronika Urbanová, Ondrej Hajdusek, Douglas F. Covey, Martin Moos, Martin Palus, Daniel Wüstner, Jan Perner

## Abstract

Sterol molecules play indispensable roles in the biology of cells and organisms. Yet, arthropods, including ticks, lack the capacity for *de novo* sterol biosynthesis and must rely on exogenous sources. How these organisms sense and utilise exogenous lipids at the cellular level remains poorly understood. Here, we show that host cholesterol and linoleic acid act as essential external signals that license proliferation of *Ixodes ricinus* tick cells. In lipid-depleted conditions, cells fail to proliferate despite retaining intracellular cholesterol, indicating that exogenous lipids function as regulatory cues rather than merely structural components. We further demonstrate that tick cells exhibit a non-canonical intracellular distribution of cholesterol, with predominant accumulation in lysosome-associated compartments, and that lipid availability induces a coordinated transcriptional program involving lipid uptake and trafficking machinery. Importantly, we find that limiting host lipid availability significantly reduces extracellular RNA output of tick-borne encephalitis virus (TBEV) from infected tick cells, indicating that viral production is affected by the metabolic state of the host cell. Together, our findings establish host-derived lipids as key regulators of cellular state in a sterol-auxotrophic organism and identify lipid availability as a determinant of flavivirus output. This work provides a conceptual framework linking nutrient sensing, cell proliferation, and vector-pathogen interactions.

## 1. Introduction

Most eukaryotes use sterols as modulators of plasma membrane properties, such as fluidity, permeability and bending rigidity ^1^. In vertebrates, sterols are acquired from the diet as well as through endogenous sterol biosynthesis. In invertebrates, the genetic make-up for sterol biosynthetic pathways has been widely lost ^2^. This genomic signature has been documented across multiple invertebrate lineages^2^, including parasitic arthropods^3,4^ and helminths^3,5^. Despite this genetic deficiency, invertebrates do sense, acquire, and utilise exogenous sterols for the regulation of key cellular and developmental processes, as well as precursors of steroid hormones ^6^. Exogenous lipids such as sterols and fatty acids are thus not only passive molecular building blocks, but also active metabolic substrates that feed multiple pathways and exert direct molecular effects on the organismal physiology and development ^7^. Ticks are obligate blood-feeding arachnids whose genomes lack the enzymatic steps to convert farnesyl pyrophosphate into sterols^8–10^, yet cholesterol is an integral component of tick physiology. How tick cells selectively acquire host cholesterol and allocate it among membranes and storage compartments remains poorly defined.

In *Drosophila*, for example, the postembryonic development does not progress on a sterol-deficient yeast diet ^11^ or on lipid-depleted medium, unless cholesterol is added back into the media ^12^, underscoring sterol essentiality for the flies’ growth and development. Lipidomics across *Drosophila* developmental stages identifies sterol esters enriched in early embryos ^13^, indicating that they could be critical for embryonic and/or postembryonic development. Beyond sterols, fatty acids such as linoleic acid also represent critical dietary components in *Drosophila*, shaping immune signalling cascades ^14^. Despite this organismal centrality, how these exogenous lipids support cell-autonomous functions in auxotrophic arthropods remains largely unexplored.

Ticks are obligate blood-feeding arthropods that transmit a wide range of pathogens of medical and veterinary importance, including tick-borne encephalitis virus (TBEV), a neurotropic flavivirus responsible for severe disease across Europe and Asia. Despite the availability of vaccines, no specific antiviral treatment exists for TBEV infection, and clinical management remains supportive. A defining feature of tick biology is their inability to synthesise sterols *de novo*, rendering them entirely dependent on host-derived lipids acquired during blood feeding. While this metabolic constraint has been recognised at the genomic level, how tick cells sense, internalise, and utilize exogenous lipids to regulate cellular processes, and whether this metabolic state influences pathogen interactions, remains poorly understood. Original studies discovered that the addition of bovine high-density lipoprotein (HDL; α-lipoprotein fraction^15^) to culture medium enhances proliferation of tick cells ^16^. However, the identity of the lipid species that drive these processes, the intracellular pathways that handle them, and their broader biological consequences remain largely unexplored in tick systems.

To address this gap, we used the *Ixodes ricinus* embryo-derived tick cell line IRE/CTVM19 ^17^ to investigate the biological impact of exogenous cholesterol and fatty acids on the biology of these cells, when cultured with delipidated serum supplementation. Here, we report that cholesterol and linoleic acid stimulate proliferation of tick cells. We further characterise the sterol specificity for the uptake and subcellular distribution of cholesterol, and identify transcriptional responses to lipid availability. Given that host lipids are known to modulate viral infection ^18^, and specifically that the low-density lipoprotein receptor (LDLR) represent a candidate entry gateway for TBEV ^19,20^ in mammals ^21^, we also examined the impact of lipid-depleted culture conditions and LDLR expression on TBEV propagation in tick cells. Collectively, this work establishes a framework for understanding the regulation and biological influence of exogenous lipids in tick biology and their role in vector-pathogen interactions.

## 2. Results

### 2.1. Exogenous cholesterol, but not other sterols, is required to sustain proliferation of sterol-auxotrophic tick cells

Similar to insects and nematodes, ticks have lost the genes required for cholesterol biosynthesis, lacking the enzymatic machinery necessary to convert farnesyl pyrophosphate into sterols (**Fig. 1A**, Supplementary Table S1). Despite lacking the biochemical capacity for *de novo* cholesterol synthesis, cholesterol was detected in all developmental stages of *I. ricinus* ticks, as well as in the embryo-derived IRE/CTVM19 cell line (**Fig. 1B, I**), suggesting that host-derived cholesterol is incorporated into tick cells. To test whether exogenous cholesterol is functionally required for tick cells, we cultured IRE/CTVM19 cells in medium supplemented with either full human serum (FS) or lipoprotein-deficient human serum (LPDS), with or without cholesterol supplementation. Cells cultured in FS showed robust multiplication, with a proliferative capacity over 5 days of a 1.51-fold increase in cell number; this was comparable to the mean 1.67-fold increase seen in cells cultured with fetal bovine serum (FBS). In contrast, cells cultured with LPDS supplementation failed to proliferate, with cell numbers remaining close to seeding density (**Fig. 1C**). However, co-supplementation of the LPDS with cholesterol conjugated to bovine serum albumin (BSA) restored proliferation, whereas BSA alone had no effect, confirming that the proliferative response was specific to cholesterol (**Fig. 1C**).

**Figure 1.**
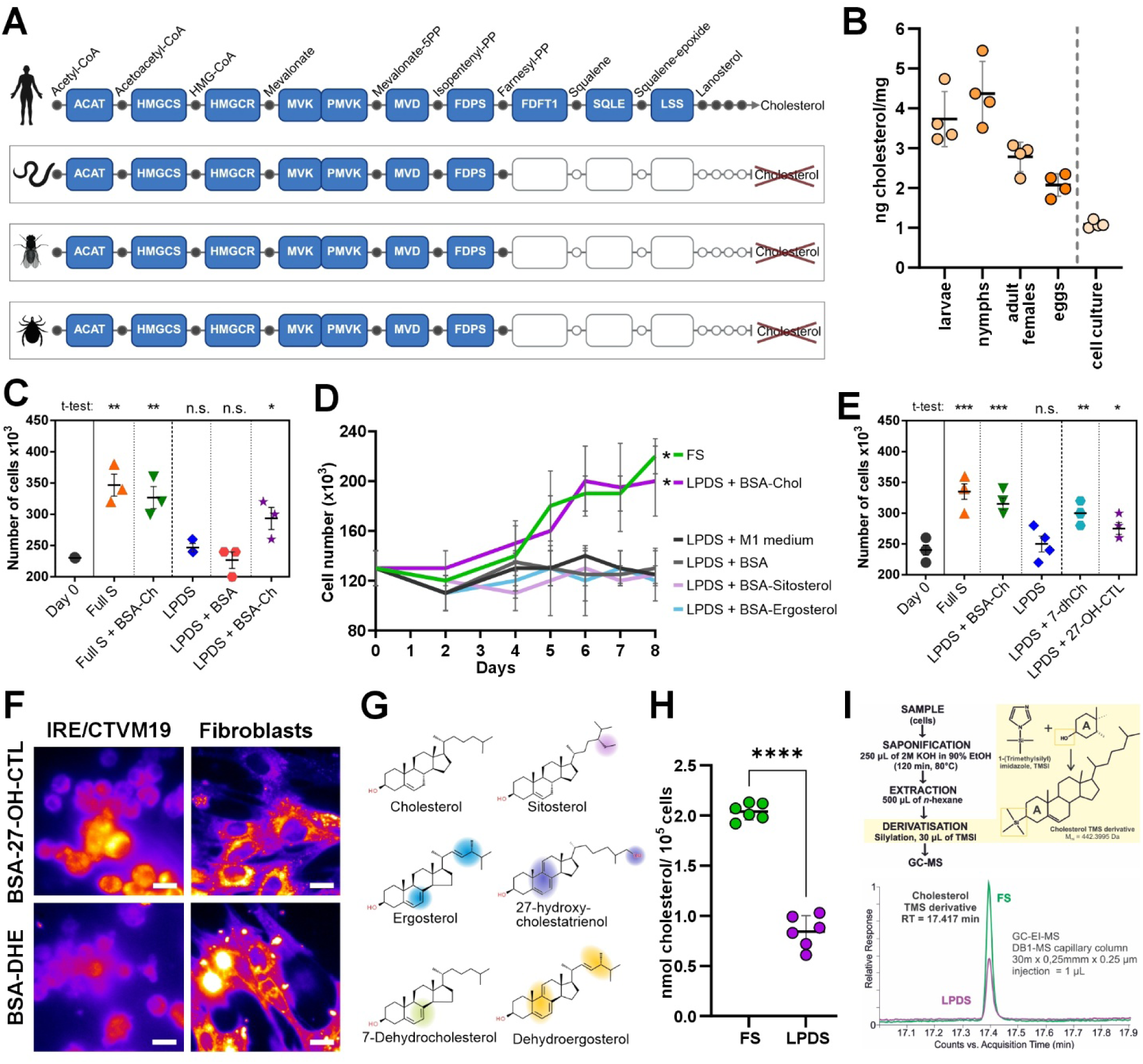
Host-derived cholesterol is required to sustain proliferation of sterol-auxotrophic tick cells. **A)** Schematic overview of the mevalonate and cholesterol biosynthesis pathway highlighting the absence of enzymatic steps required for de novo cholesterol biosynthesis in *Ixodes ricinus* ticks, *Drosophila*, and nematodes (see Supplementary Table 1). Metabolic intermediates are indicated with circles, and enzymatic steps are shown with rectangles using standard abbreviations: ACAT, acetyl-CoA C-acetyltransferase; HMGCS, 3-hydroxy-3-methylglutaryl-CoA synthase; HMGCR, 3-hydroxy-3-methylglutaryl-CoA reductase (NADPH-dependent); MVK, mevalonate kinase; PMVK, phosphomevalonate kinase; MVD, diphosphomevalonate decarboxylase; FDPS, farnesyl diphosphate synthase; FDFT1, farnesyl-diphosphate farnesyltransferase (squalene synthase); SQLE, squalene monooxygenase; LSS, lanosterol synthase. The terminal 12 enzymatic steps converting lanosterol to cholesterol are not shown. **B)** Quantification of cholesterol content of unfed *I. ricinus* developmental stages and IRE/CTVM19 tick cells. Cholesterol is expressed as ng per mg of dry weight for ticks and per mg of cell pellet for cultured cells. **C)** The effect of exogenous cholesterol on IRE/CTVM19 cell proliferation after 5 days of culture. Cells were grown in medium supplemented with either full human serum (FS) or lipoprotein-deficient serum (LPDS), both supplemented with cholesterol (Chol) conjugated on BSA as a vehicle (Supplementary Fig. S1). Cell numbers were determined per 1 mL of culture medium. **D)** Growth curves of IRE/CTVM19 cells cultured for 8 days in FS or LPDS supplemented with various sterols. Supplements included BSA-conjugated cholesterol, sitosterol, or ergosterol. LPDS supplemented with M1 medium and LPDS with BSA served as negative controls. Data are shown as mean ± SD. **E)** Effect of modified sterols on IRE/CTVM19 cell proliferation. Cells were cultured in LPDS supplemented with BSA-conjugated cholesterol, BSA-conjugated 7-dehydrocholesterol (BSA-7-dhCh), or BSA-conjugated 27-hydroxy-cholestatrienol (BSA-27-OH-CTL). Cell counts were determined after 5 days and from 1 mL of culture medium. **F)** Uptake of fluorescent sterol analogues visualised by epifluorescence microscopy in tick cells and mammalian fibroblasts. Cells were incubated in LPDS-supplemented medium with BSA-conjugated 27-hydroxy-cholestatrienol (BSA-27-OH-CTL) or BSA-conjugated dehydrocholesterol (BSA-DHE). UV-range fluorescence signal indicates intracellular sterol accumulation. Control cells received medium only. Representative images are shown. Control images are shown in Supplementary Fig. S2. **G)** Structural differences between cholesterol and selected sterol analogues. **H)** Cellular cholesterol content of IRE/CTVM19 cells cultured in full serum (FS) or LPDS, expressed as nmol cholesterol per 10⁵ cells. Individual points represent biological replicates. **I)** Workflow for cholesterol quantification of tick tissues and IRE/CTVM19 tick cells by gas chromatography–mass spectrometry (GC-MS). Samples were subjected to saponification, extraction and derivatisation using 1-(trimethylsilyl)imidazole (TMSI) to generate cholesterol trimethylsilyl (TMS) derivative (monoisotopic mass, Mmi = 442.3995 Da). A representative GC-MS chromatogram of cholesterol TMS derivative is shown (retention time = 17.417). Data in graphs are presented as mean ± SEM. Significance was determined using a t-test: *P* < 0.05 (*), *P* < 0.01 (**), *P* < 0.001 (***), *P* < 0.0001 (****), n.s. = not significant.

To assess the structural specificity of sterols inducing cell proliferation, we tested structurally-related sterols. Sitosterol and ergosterol, which share the sterol backbone with cholesterol but differ in side-chain structure and double bond configuration, were unable to rescue proliferation with LPDS supplementation even when assessed over an 8-day period (equivalent to the estimated doubling time of FS-cultured cells) (**Fig. 1D**). To further investigate the relationship between sterol structure and function, we next examined sterols with modifications to the ring system and side chain, focusing on compounds that also possess intrinsic fluorescence to enable tracking of uptake. IRE/CTVM19 cells were cultured in LPDS supplemented with either BSA-conjugated 7-dehydrocholesterol (DHE) or BSA-conjugated 27-hydroxy-cholestatrienol (27-OH-CTL), and cell proliferation was assessed after 8 days (**Fig. 1E, G**). These two sterols partially rescued cellular proliferation (**Fig. 1E**). These findings suggest that side chain and ring modifications strongly influence uptake and proliferative rescue in tick cells.

To further investigate sterol selectivity, we examined the uptake of intrinsically fluorescent sterol analogues using UV-specialised wide-field microscopy. Tick cells efficiently internalised 27-OH-CTL, but not DHE, when both were delivered as BSA-conjugates (**Fig. 1F, G**, Supplementary Fig. S2). This selective uptake supports our earlier findings, indicating that bulky side-chain modifications (such as those found in ergosterol) impair uptake by tick cells and thereby limit their ability to stimulate proliferation. In contrast, human fibroblasts readily internalised DHE (**Fig. 1F,G**; Supplementary Fig. S2), highlighting a fundamental difference in sterol uptake specificity between ticks and mammals). Using the GC-MS quantification, we further demonstrated that FS-grown cells accumulated significantly more cholesterol than LPDS-grown cells (*P* < 0.0001) (**Fig. 1H, I**), with cells in FS containing 2.038 ± 0.034 nmol of cholesterol / 10^5^ cells (mean ± SEM, n = 6) and those in LPDS containing 0.843 ± 0.065 nmol of total cholesterol / 10^5^ cells (mean ± SEM, n = 6). The fact that cells proliferate only with higher cholesterol amounts suggests that proliferation correlates with a higher intracellular cholesterol pool. Together, these results demonstrate that cholesterol is a critical exogenous factor required for tick cell proliferation and underscore the structural specificity and selectivity of the tick sterol uptake machinery.

### 2.2. Tick cells display predominant intracellular cholesterol localisation compared to plasma membrane enrichment in mammalian cells

To investigate how cholesterol is distributed within tick cells, we stained *I. ricinus* IRE/CTVM19 cells with filipin, which binds to unesterified cholesterol, and with the plasma membrane co-stain CellBrite Fix 640. In contrast to mammalian cells (Vero E6), which showcase the characteristic localisation of cholesterol to the plasma membrane, tick cells displayed strong filipin signals within intracellular regions (excluding the nucleus), with only a limited colocalisation with the plasma membrane, displaying plasma membrane-to-cytoplasm filipin intensity ratios below 1 (**Fig. 2A, B, C**). Filipin staining showed a similar ratio of intracellular-to-plasma membrane deposition of cholesterol in both proliferating (FS; 0.716 ± 0.036, mean ± SD) and non-proliferating (LPDS; 0.654 ± 0.049, mean ± SD) tick cells (**Fig 2C**), indicating that lipid starvation does not induce major reallocation of intracellular cholesterol pools. Presence of chloroquine (Chlq), a lysosomotropic agent that disrupts endo-lysosomal acidification^22^, or U18666A (U18), an inhibitor of cholesterol export from lysosomes that interferes with the function of Niemann-Pick 1 (NPC1) protein^23^, caused significantly decreased depositions of cholesterol in the plasma membrane of tick cells (**Fig. 2C)**. Consistent with a lysosome-dependent cholesterol trafficking step, the two inhibitors of lysosomal function, chloroquine and U18666A, directly impaired cell proliferation in a dose-dependent manner (Supplementary Fig. S3). Together, these data support a model in which lysosomes act as an essential intermediate between cholesterol internalization and downstream trafficking, including delivery to the plasma membrane.

**Figure 2.**
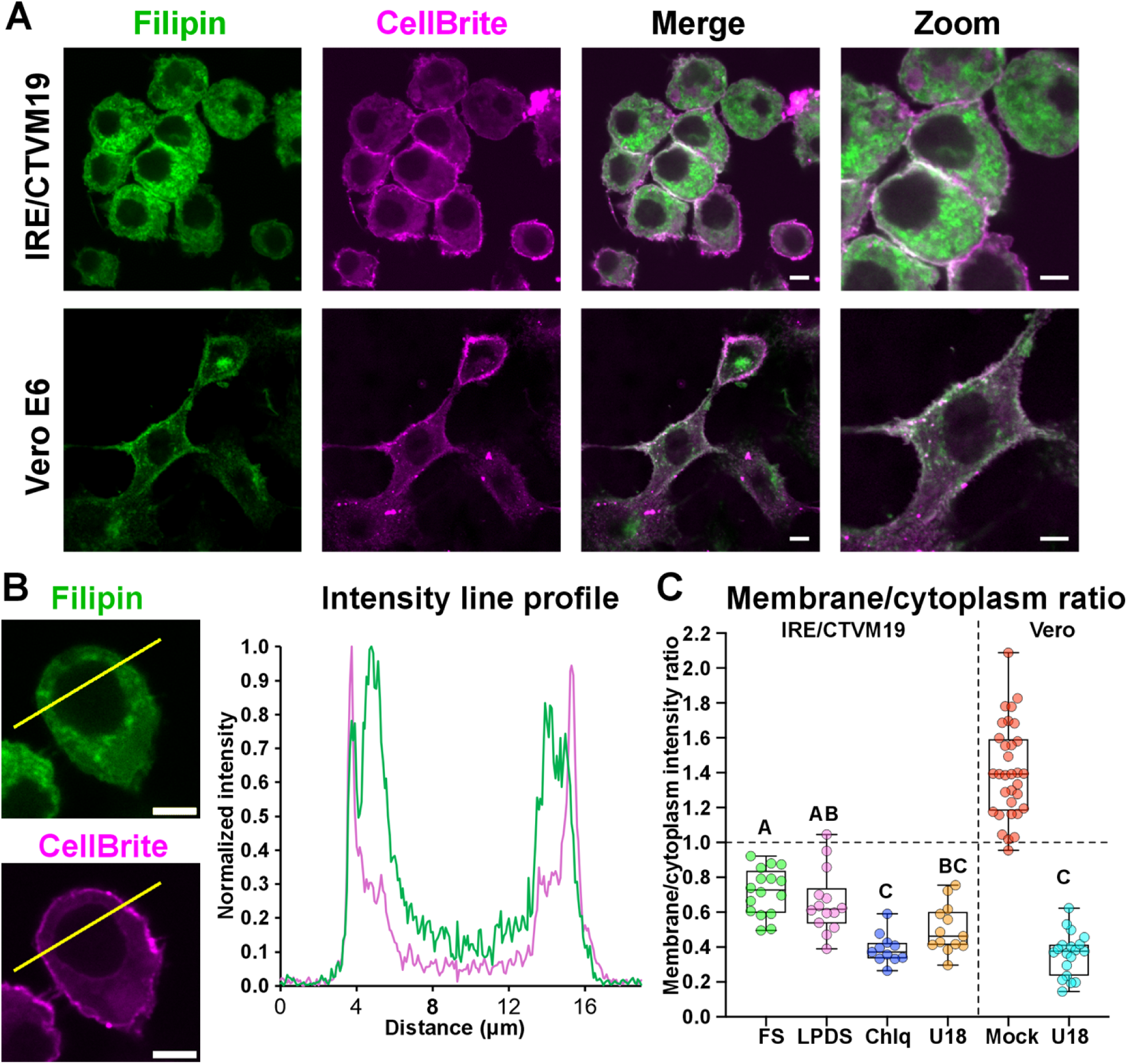
Tick cells exhibit a non-canonical intracellular distribution of cholesterol. **A)** Representative confocal images of unesterified cholesterol in IRE/CTVM19 tick cells and Vero E6 mammalian cells. Cells were stained with filipin (green) to label free cholesterol and with CellBrite Fix 640 (magenta) to mark the plasma membrane. Merged images highlight the spatial distribution of cholesterol relative to the membrane. White color in the merged images indicates overlap of filipin and membrane staining. **B)** Representative line intensity profiles across a single IRE/CTVM19 cell showing normalized filipin and CellBrite signal intensity. The intracellular predominance of filipin signal is evident compared to limited overlap with the membrane marker on the cell periphery. Scale bars in all images: 5 μm. **C)** Quantification of filipin signal distribution between the plasma membrane and intracellular compartments (excluding the nucleus) in IRE/CTVM19 cells under various conditions. Membrane-to-cytoplasm intensity ratios were measured for cells cultured in full serum (FS), lipoprotein-deficient serum (LPDS), or FS co-supplemented with chloroquine (Chlq) or U18666A (U18). Vero E6 cells were included as a control and displayed higher filipin intensity at the plasma membrane. Data in graphs are presented as mean ± SD. Different letters indicate statistically significant differences between groups (one-way ANOVA with post hoc test). Groups sharing at least one letter are not significantly different from each other (*P* < 0.05). Images used for line intensity analysis are shown as Supplementary Fig. S4.

### 2.3. Tick cells localise part of intracellular cholesterol to lysosomes and lipid droplets

To test this notion and determine the specific intracellular distribution of cholesterol deposits in tick cells, we performed co-localisation studies using filipin and either LD540 or LysoTracker Red (**Fig. 3A**). While LysoTracker is a dye very specific for lysosomes, LD540 is a lipophilic dye, which partitions into lipid droplets and is therefore used as marker for these organelles within cells. The line profile analyses showed that filipin partially overlaps with LysoTracker signals, and also partially with LD540 (**Fig. 3B**). Using a semi-automatic image segmentation protocol, we analysed the fluorescent co-stains quantitatively. The Pearson’s correlation coefficients (*r*) support a positive correlation between filipin (marker for cholesterol) and LysoTracker signals (marker for lysosomes) and, to a slightly lower extent, a positive correlation between filipin and LD540 (marker for lipid droplets), indicating that cholesterol is largely localised to these compartments in tick cells (**Fig. 3C**). Altogether, these data demonstrate that IRE/CTVM19 tick cells accumulate a fraction of free, i.e. non-esterified, cholesterol localised in lysosomes and lipid droplets. However, the median *r* values <0.4 indicate that a substantial amount of intracellular cholesterol remains unassigned to a cellular destination, likely associated with other subcellular compartments in tick cells.

**Figure 3.**
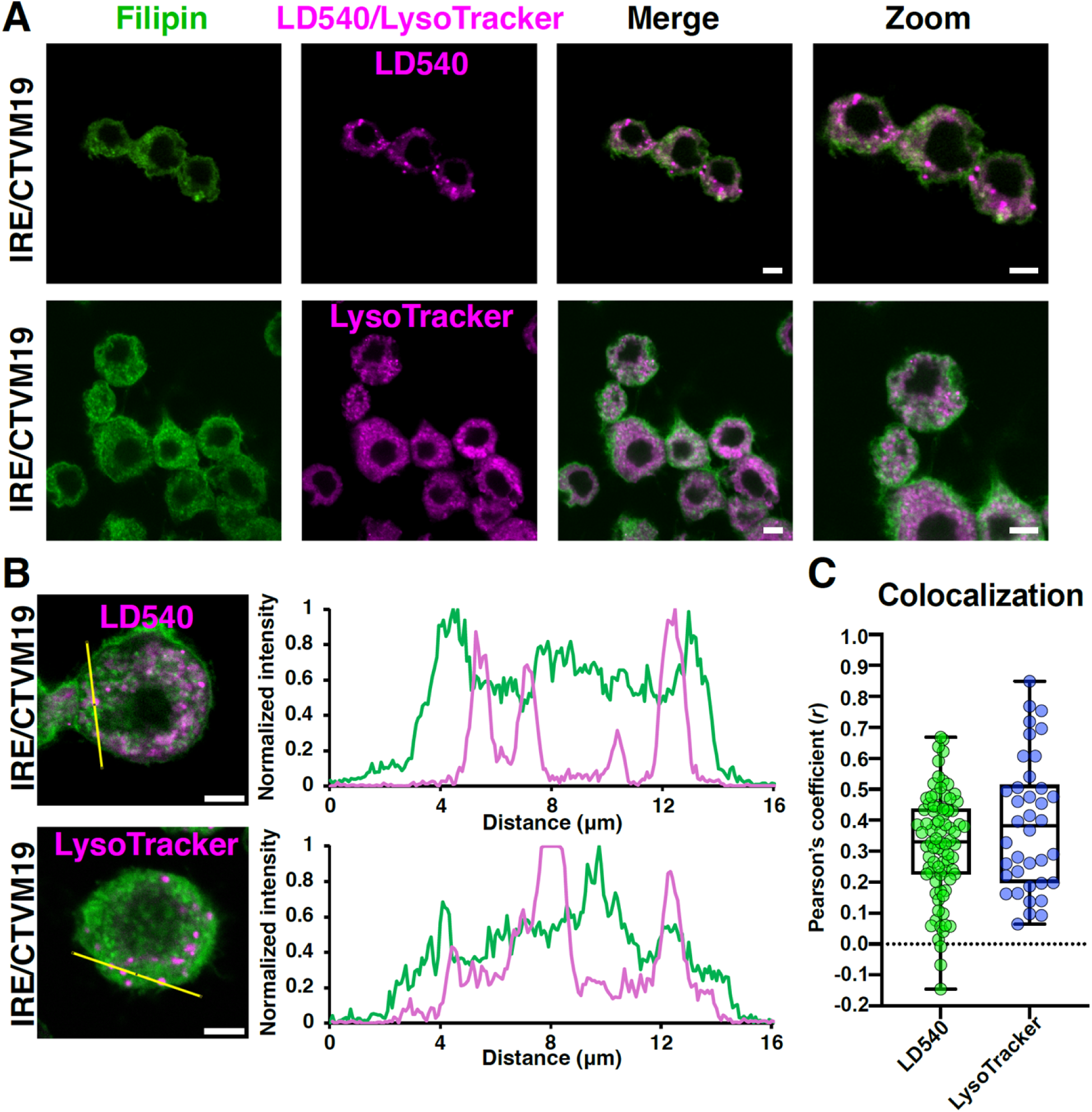
Intracellular cholesterol localises to lysosomes and lipid droplets in tick cells. **A)** Localisation of filipin (marker for cholesterol) and LD540 or LysoTracker Red which stain for lipid droplets and lysosomes, respectively, in IRE/CTVM19 cells. **B)** Representative line profiles across IRE/CTVM19 cells stained with filipin and LD540 or LysoTracker Red. Partial accumulation of filipin in LD540-stained and LysoTracker-stained regions is discernible. **C)** Quantification of filipin colocalisation with LD540-stained and LysoTracker-stained regions in IRE/CTVM19 cells. Each data point represents a colocalisation measurement for a single cell. The whole cell (minus nucleus) is used for colocalisation determination in this analysis, with 82 cells analysed for LD540 and 38 cells for Lysotracker.

### 2.4. Linoleic acid acts as an additional lipid signal that licenses tick cell proliferation

To assess whether fatty acids are essential for tick cell proliferation, IRE/CTVM19 cells were cultured in LPDS supplemented with individual BSA-conjugated fatty acids (**Fig. 4A**). As expected, for cells cultured in full serum (FS) the cell number increased within five days again by about 50%, whereas cells cultured in LPDS failed to proliferate and remained close to seeding density (**Fig. 4B**). Supplementation with BSA carrier alone also did not promote proliferation, confirming that the BSA alone had no proliferative effect when used as a fatty acid-vehicle. Among the fatty acids tested, linoleic acid (18:2n-6) significantly enhanced proliferation, with cell counts reaching levels comparable to FS (*P* < 0.05). In contrast, supplementation with palmitate (16:0), oleate (18:1n-9), γ-linolenate (18:3n-3), or arachidonate (20:4n-6) did not significantly increase cell numbers compared to LPDS controls (all *P* ≥ 0.05). To test for an additive or synergistic effect of linoleic acid and cholesterol, we analysed growth capacity of tick cells in LPDS media supplemented with BSA-cholesterol and BSA-linoleic acid separately and together. The data showed that each of these supplementations sustained tick proliferation to a similar extent, with no additional effect of BSA-cholesterol and BSA-linoleic acid co-supplementation (Supplementary Fig. S5). Consequently, both exogenous lipid species, cholesterol and fatty acids, are likely required for the same cellular function. Thus, linoleic acid emerges as a unique exogenous fatty acid capable of sustaining tick cell proliferation in the absence of serum-derived lipids. These findings identify linoleic acid, alongside cholesterol, as a critical exogenous lipid input for tick cells, suggesting that tick cells can sense and utilise linoleic acid as a lipid signal to support proliferation.

**Figure 4.**
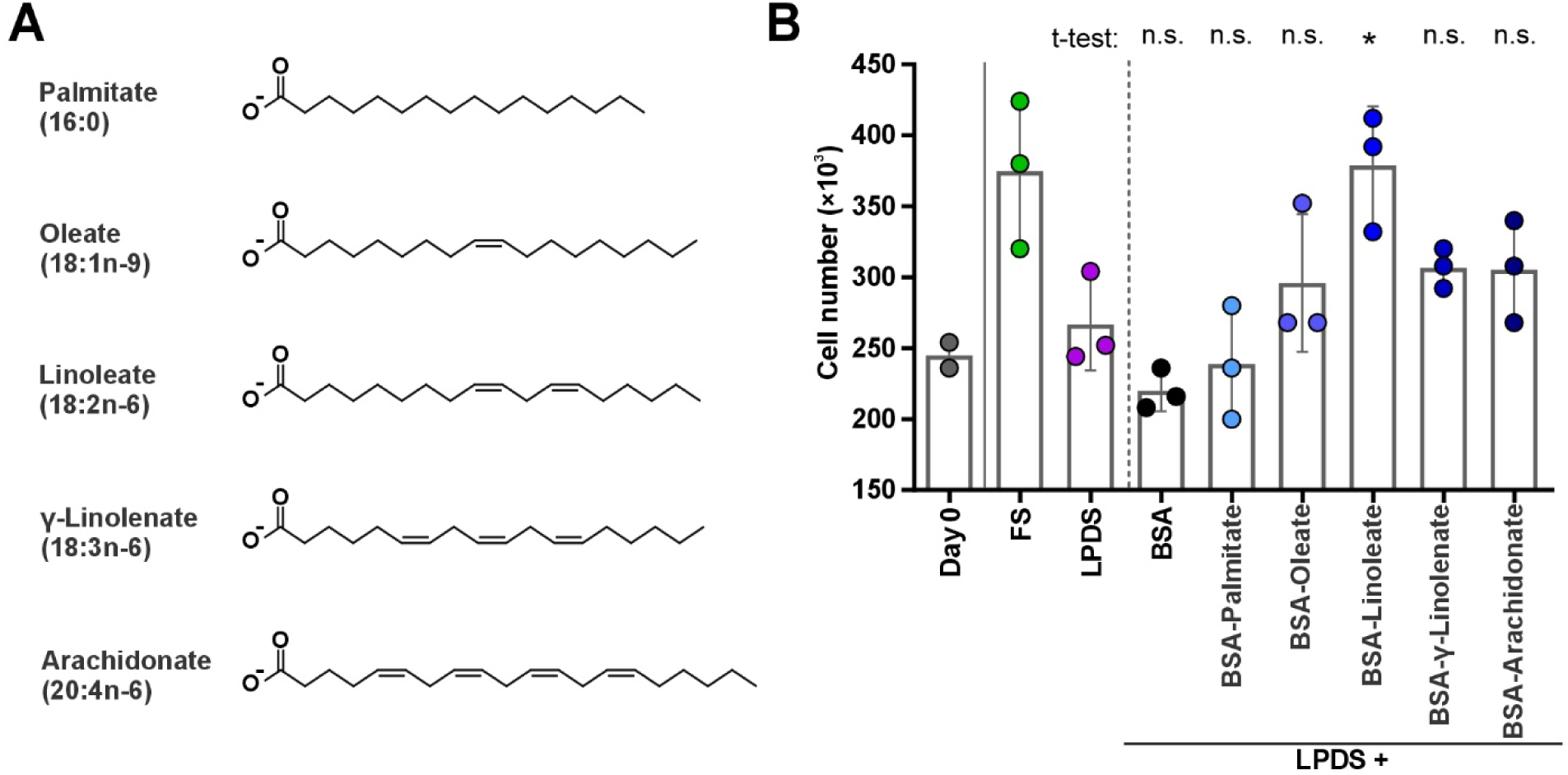
Linoleic acid acts as a lipid signal that supports proliferation of tick cells. **A)** Structural formulae of the fatty acid salts used for BSA conjugation and supplementation of LPDS medium: palmitate (16:0), oleate (18:1n-9), linoleate (18:2n-6), γ-linolenate (18:3n-6), and arachidonate (20:4n-6). **B)** IRE/CTVM19 cells were cultured for 5 days in medium supplemented with full serum (FS) or lipoprotein-deficient serum (LPDS). LPDS cultures were supplemented with BSA alone (vehicle control) or BSA-conjugated fatty acid salts (palmitate 16:0, oleate 18:1n-9, linoleate 18:2n-6, γ-linolenate 18:3n-6, arachidonate 20:4n-6) at 15 µM. Cell numbers were determined as counts per 1 mL culture medium. Each dot represents an independent culture (n = 3); bars show mean ± SD. Statistical significance was assessed by unpaired t-test relative to LPDS controls; only BSA-linoleate significantly increased cell numbers (*P* < 0.05).

### 2.5. Exogenous lipids induce a transcriptional program associated with lipid uptake and lysosomal trafficking

To investigate whether lipid cues influence expression of genes involved in lysosomal handling of cholesterol, we quantified transcripts encoding Niemann-Pick proteins (NPC1, NPC2 variants) and the lysosomal integral membrane protein-2 (LIMP-2) in IRE/CTVM19 cells cultured for 7 days. While *npcs* transcripts were found to be mostly unregulated by exogenous lipids, with the exception of *npc2-2* (**Fig. 5A, B**), levels of *limp* transcripts were significantly elevated in the presence of exogenous lipids at later time-points of culturing (**Fig. 5C**), which correlate with the proliferative phase. LIMP-2 belongs to the CD36 family of scavenger receptors, a member of which, named Croquemort, was recently shown to be pulled down by 1-palmitoyl-2-oleoyl-sn-glycero-phosphoglycerol beads from an *Ixodes scapularis* cell line lysate, but showed localisation to the plasma membrane ^24^. As *I. ricinus* LIMP-2 shares the protein 3D architecture (Supplementary Fig. S6), as well as primary amino acid sequences (Supplementary Fig. S7), with *I. scapularis* Croquemort, we wanted to experimentally verify the cellular targeting of *I. ricinus* LIMP-2. To confirm the predicted lysosomal targeting of tick LIMP-2, we used an established *Drosophila* S2 expression system (Thermo Fisher Scientific) to produce recombinant *I. ricinus* LIMP-2 protein (**Fig. 5D**), coupled with a C-terminal mCherry fluorescent reporter. We showed a clear co-localisation of tick LIMP-2 with LysoTracker in S2 cells (**Fig. 5E**), confirming the localisation of tick LIMP-2 homologue to lysosomes. Given the accumulation of cholesterol in lysosomes under full serum conditions, as indicated by filipin staining, the upregulation of LIMP-2 may reflect its role in lysosomal cholesterol handling.

**Figure 5.**
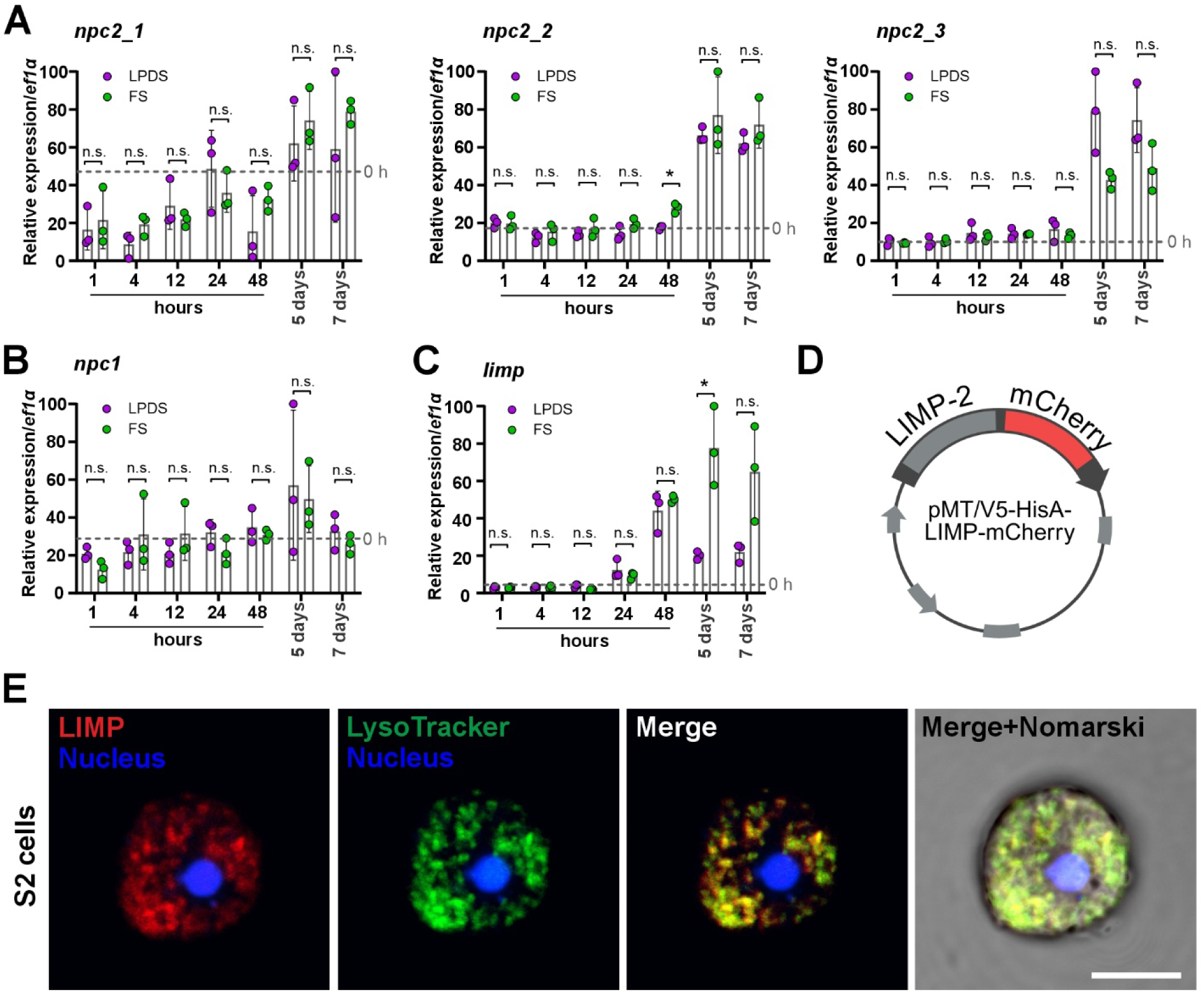
Lipid availability induces transcriptional responses associated with lysosomal cholesterol handling. A-C) Relative expression (RT-qPCR) of transcripts encoding proteins implicated in cholesterol efflux from lysosomes in the IRE/CTVM19 cells throughout 7 days of culturing with full serum (FS) or delipidated serum (LPDS) supplementation. Each dot represents an independent culture. T test values: * *P* < 0.05; n.s. = not significant *P* ≥ 0.05. npc2_1: Niemann Pick 2 from 1, IrSigP-112437; npc2_2: Niemann Pick 2 from 2, IrSigP-112437; npc2_3: Niemann Pick 2 from 3, IrSigP-112437; npc1, Niemann Pick 1, ID: Ir-110144; limp-2, lysosomal integral membrane protein, gene ID: IricT00008008-PA. Expression values of targeted genes were normalised to expression values of *elongation factor 1 alpha*. **D)** A schematic of construct for *Drosophila* S2 cells heterologous expression of *I. ricinus* LIMP-2-mCherry. **E)** *Drosophila* S2 cells expressing *I. ricinus* LIMP fused to mCherry (red) were stained with LysoTracker (green) and Hoechst (blue). Confocal imaging shows co-localization of LIMP with lysosomes. Scale bar: 5 μm

Finally, to reveal a full program of transcriptomic regulation, responsive specifically to external cholesterol and linoleic acid, we performed RNA-sequencing of IRE/CTVM19 cells subjected to differential availability of exogenous lipids (Supplementary Fig. S8). Specifically, we compared cells grown with *i*) FS versus LPDS supplementation, *ii*) LPDS + BSA-cholesterol versus LPDS + BSA control, or *iii*) LPDS + BSA-linoleate versus LPDS + BSA control. Cells were either not pre-starved (nps) and incubated for five days in differential media and compared as above (*i–iii*), or pre-starved (ps) in LPDS for five days and only then cultured for two days or six days in differential media and then compared as above (*i–iii*) (Supplementary Fig. S8). Acquired Illumina reads were assembled with a high percentage of BUSCO completeness (Supplementary Fig. S9), indicating robust transcriptome assembly quality and gene content coverage. We identified a broad transcriptomic change in response to available lipids. Notably, these complex expression signatures regulated in response to lipid cues include *srebp* and *fatty acid synthase*, both induced by all three treatments: supplementation with full serum, or specifically with cholesterol, or specifically with linoleic acid (Supplementary Fig S10). Together, these findings point to a broad lipid-responsive reprogramming of cellular gene expression governing the proliferation of tick cells.

### 2.6. Lipid deficiency increases LDL receptor-family expression and constrains TBEV RNA output

To investigate whether extracellular lipoprotein availability influences biology of TBEV, we cultured IRE/CTVM19 cells in medium supplemented with full human serum (FS) or lipoprotein-deficient serum (LPDS) and infected them with TBEV (**Fig. 6A**). Commercial human sera from a non-TBEV-endemic region were used to minimise the likelihood of pre-existing anti-TBEV antibodies as a confounding factor. Recent studies identified the low density lipoprotein receptor (LDLR) as a possible entry receptor for TBEV in mammals ^19,20^. We searched the *I. ricinus* genome for related proteins and identified a candidate LDL receptor-family transcript, IricT00002028 (**Fig. 6B**). We next measured expression of this transcript in IRE/CTVM19 cells cultured under full-serum or LPDS conditions and revealed its higher abundance in LPDS than in full-serum cultures after 7 days (**Fig. 6C**). To determine whether enhanced LDLR expression was associated with increased viral entry into the cell, tick cells were pre-starved for 9 days in LPDS, then infected with TBEV, and then infected cells were maintained under FS or LPDS conditions for 3 days. Persistent LPDS culture did not significantly increase levels of intracellular TBEV RNA, normalised to tick *ef1α* (*P* = 0.1; **Fig. 6D**). In contrast, extracellular TBEV RNA copy numbers in the supernatant were significantly lower TBEV RNA in LPDS than in full-serum cultures (**Fig. 6E**). These data indicate that cellular lipid homeostasis influences extracellular TBEV RNA production in IRE/CTVM19 cultures.

**Figure 6.**
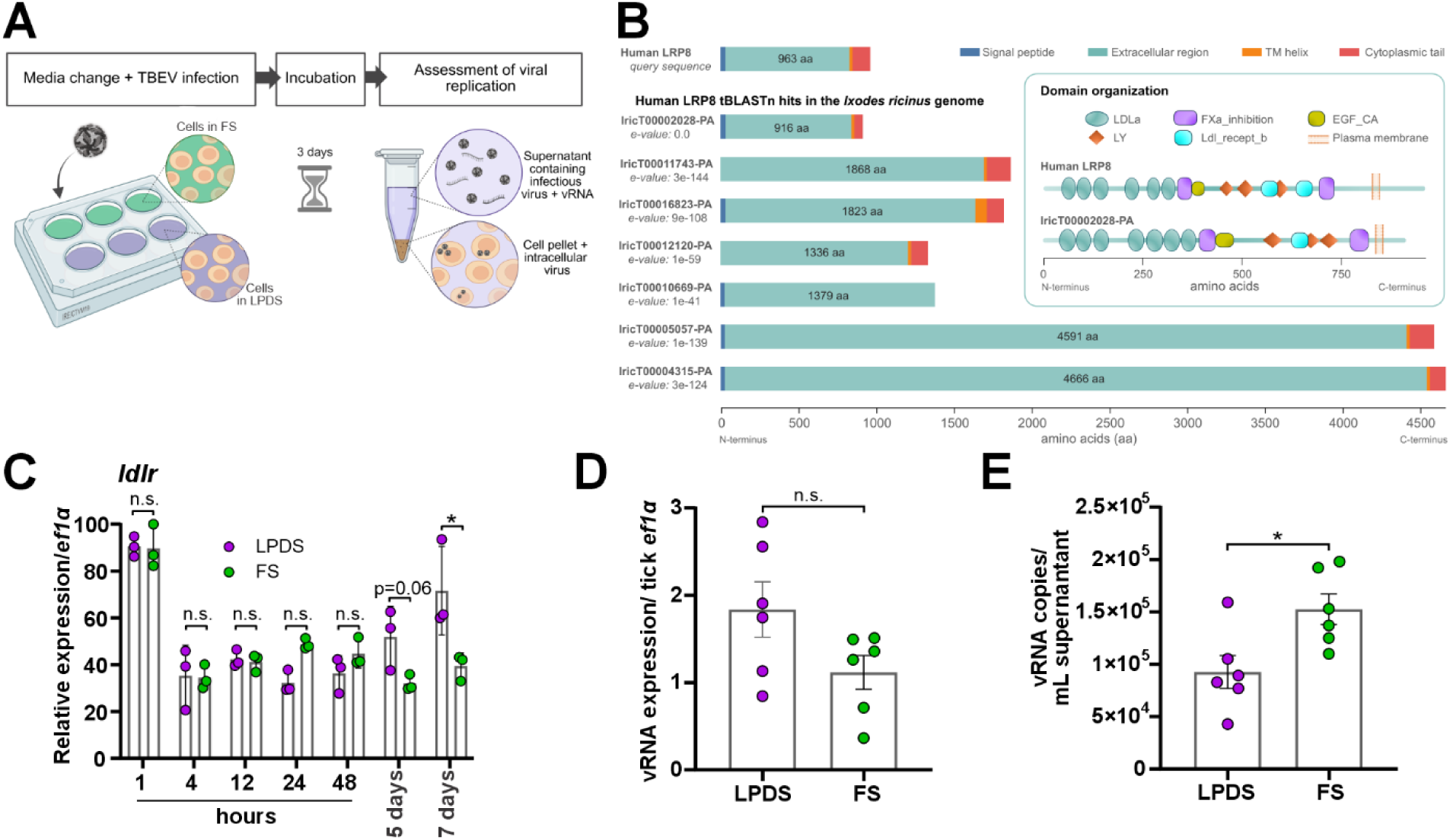
Lipid availability modulates LDL receptor expression and affects TBEV RNA output in tick cells. **A)** Schematic overview of the experimental design. IRE/CTVM19 cells were infected with TBEV and maintained in medium supplemented with full human serum (FS) or human lipoprotein-deficient serum (LPDS). At 3 days post-infection, culture supernatants and cell pellets were collected for viral RNA quantification by RT-qPCR. **B)** Homology-based identification of human LRP8/LDL receptor-family protein hits in the *Ixodes ricinus* genome. Predicted protein lengths, e-values, signal peptides, extracellular regions, transmembrane helices, cytoplasmic tails, and selected conserved domains are shown. The candidate transcript used for expression analysis is indicated as IricT00002028. **C)** Relative expression of the *I. ricinus* LDL receptor-family transcript (*ldlr*; IricT00002028) in IRE/CTVM19 cells cultured in FS or LPDS for the indicated times. Transcript abundance was quantified by RT-qPCR and normalised to tick *elongation factor 1α* (*ef1α*). **D)** Cell-associated TBEV RNA abundance at 3 days post-infection, quantified by RT-qPCR and normalised to tick *ef1α* mRNA. **E)** TBEV RNA copy number in culture supernatants at 3 days post-infection, quantified by RT-qPCR and expressed as viral RNA copies per mL. In C–E, each dot represents an independent culture; bars show mean ± SEM. Statistical significance was assessed by t-test (C), or Mann Whitney (D,E). *P* < 0.05; n.s., not significant.

## 3. Discussion

Our work provides the cell-level demonstration that exogenous cholesterol and linoleic acid together sustain tick cell proliferation, likely via activation of a lipid-sensing regulatory pathway. The loss of endogenous sterol biosynthesis in ticks is in line with the genome history of other invertebrates^2,5^, with sterol auxotrophy being a broader phenomenon among them. This isoprenoid intermediate is instead, in the absence of downstream sterol-forming enzymes, expected to be diverted into essential non-sterol products, including protein prenylation, the polyprenyl side chain of ubiquinone and dolichol, and farnesylated haem *o*/haem *a* ^25^. A fraction of this pool could, in principle, feed into methyl-farnesoate/juvenile-hormone-like products, but tick genomes lack clear orthologues of key juvenile-hormone branch enzymes^8^ and direct biochemical evidence for methyl-farnesoate synthesis in ticks remains limited ^26^.

Species of at least some invertebrate lineages have retained functional dependency on sterol molecules and are thus dependent on dietary acquisition ^5^. Exogenous sterols also represent molecules indispensable in physiology and development of ticks, with the most studied role being host cholesterol as a substrate for steroid hormone synthesis. Exogenous ecdysteroid administration has shown that tick development can be experimentally induced by exogenous 20-hydroxyecdysone (20E), triggering early vitellogenin expression and oogenesis in virgin females ^27–29^. When supplied generously suspended in the blood meal, ecdysterone can even provoke aberrant “supermolting” of adult ticks ^30^. These observations underscore the absolute reliance of ticks on acquired exogenous sterols, as endocrine substrates, and highlight the need to resolve how individual tick cells internalise, traffic, and allocate sterols to sustain growth and reproduction. Here, we quantified cholesterol deposits across developmental stages of *I. ricinus* ticks and determined that ticks harbour nanogram quantities of host-derived cholesterol (per milligram of biomass) with consistent deposition across larval, nymphal, and adult stages. These values illustrate that cholesterol storage is a pervasive feature of tick development rather than restricted to a single life stage. Original studies suggested that a large proportion of dietary cholesterol is being expelled as cuticle wax^31^ or stored in pheromone glands ^32^. Comparable concentrations were also detected in cultured *I. ricinus* cells, indicating that dietary cholesterol is retained and mobilised at the cellular level. Pioneering studies that optimised culture medium composition for cell lines derived from *Rhipicephalus* (*Boophilus*) *microplus*, *Rhipicephalus appendiculatus* and *Dermacentor* (*Anocentor*) *nitens* ticks also noticed a positive proliferative effect of cholesterol (if given at 5–10 µg/mL) in the presence of FBS^16^, indicative of a role in stimulation of multiplication. In vertebrate cells, by contrast, cholesterol is an abundant bulk lipid, with macrophages containing around 100 ng cholesterol per 10⁵ cells^33^. This contrast indicates that cholesterol in ticks likely serves as a limiting substrate or a signal-like cue, rather than simply a structural lipid. This would be in line with observations in *C. elegans*, another sterol auxotroph, where sterols are required in minute amounts and appear to act largely as regulatory/hormone-like factors controlling developmental transitions rather than as bulk membrane constituents^34,35^. In organisms devoid of internal sterol biosynthesis, presence of external sterol might represent a molecular key that triggers proliferative, developmental ^36^, and reproductive ^37^ processes.

The absence of *de novo* sterol biosynthesis in ticks ^8–10,26^ raises questions regarding how these ectoparasites acquire and utilise sterols. Our findings indicate that *I. ricinus* cells have evolved specialised pathways for internalisation and subcellular trafficking of exogenous cholesterol. Uptake of host cholesterol in ticks most likely occurs *via* endocytic routes analogous to those in other animals, wherein cholesterol-rich lipoproteins (e.g. low-density lipoprotein, LDL) are engulfed and delivered to lysosomes. In mammalian systems, LDL-derived cholesteryl esters within late endosomes/lysosomes are hydrolysed to free cholesterol, which is then handed over by the soluble luminal carrier NPC2 to the N-terminal luminal domain of the multi-spanning membrane transporter NPC1 ^38,39^. NPC1 then inserts cholesterol into the limiting lysosomal membrane for onward distribution ^40^. In parallel to the canonical NPC1/NPC2 route, the lysosomal membrane protein LIMP-2 directly binds cholesterol via a luminal tunnel in its ectodomain and facilitates export of lipoprotein-derived cholesterol ^41^, possibly facilitating cholesterol channelling between lysosomes and ER ^42^. These findings position LIMP-2 as an auxiliary lysosomal cholesterol transporter that can operate alongside NPC1. In *C. elegans*, mutants of *npc1* display abnormal sterol accumulation in the body cavity^43^, indicating the key role of NPC1 in systemic distribution of cholesterol after its acquisition. In soft ticks, an NPC1-like protein was identified and its essentiality for post-feeding survival was demonstrated^44^. Here, using the NPC1 inhibitor U18666A, we reduced lysosome-plasma membrane trafficking in tick cells, validating a function of NPC1 in cholesterol export of lysosomes in these cells. Also, previous work has identified a CD-36 family member (Croquemort), which was characterised as a protein functionally and structurally related to LIMP-2, but determined its localisation to the cell membrane rather than lysosomes ^24^. Here, we identified a 1:1 homologue of mammalian LIMP-2 and experimentally verified its predicted (DeepLoc 1.0 ^45^; Likelihood: 0.6) lysosomal targeting.

The transcriptional response of tick cells to exogenous cholesterol and linoleic acid suggests the existence of a lipid-sensing program that couples extracellular lipid availability to proliferation ^46^. Conserved regulators such as the SCAP-SREBP system ^47^ or nuclear hormone receptor pathways^48^ are plausible candidates, as homologous components are encoded in tick genomes ^8^. In other metazoans, these pathways link sterol and fatty-acid availability to membrane lipid homeostasis and growth, although their sensing specificity differs between the sterol-sensing mammalian system ^49^ and the palmitate-sensing system of *Drosophila* ^50,51^. Our data further point to an important role for lysosome-associated cholesterol handling in this response, because cholesterol accumulates predominantly in intracellular compartments, and pharmacological disruption of lysosomal cholesterol export suppresses proliferation. We therefore propose that tick cells integrate exogenous sterol and fatty-acid cues through a lipid-responsive program linked to lysosomal trafficking.

Lipid deprivation induced expression of the tick LDLR, consistent with a compensatory response to reduced extracellular lipoprotein availability and raising the possibility that LPDS culture might potentiate TBEV entry. This interpretation is conceptually supported by recent studies identifying the LDLR-family member LRP8 as an entry receptor for TBEV, with receptor depletion reducing infection, and ectopic expression enhancing viral attachment and internalization in mammalian cells ^19,20^. However, we found no statistically supported evidence that the lipid-responsive increase in LDLR expression altered intracellular levels of viral RNA in tick cells. This view is consistent with the broader understanding, in which flavivirus infection depends not only on entry factors but also on lipid-dependent membrane remodeling required for genome replication and virion production ^52,53^. This interpretation is supported by broader viral systems in which cholesterol trafficking and lysosomal lipid homeostasis directly modify virus production: Respiratory syncytial virus (RSV) infection, for example, blocks cholesterol transport from lysosomes to the ER, activates the SREBP2-LDLR axis, increases LDL uptake, and promotes cholesterol-rich lysosomal dysfunction that supports RSV replication and viral protein accumulation ^54^. Thus, perturbation of lipid compartments may restrict infection in several models, and pharmacological targeting ^55^ of cholesterol, sphingolipid, fatty-acid or lipid-droplet pathways has repeatedly been proposed as an antiviral strategy, although the direction and magnitude of the effect remain virus-and cell-type-specific ^56^. In this framework, the unusual cholesterol organization of IRE/CTVM19 cells, where cholesterol is preferentially intracellular and partly associated with lysosome-like compartments and lipid droplets, suggests that tick cells may impose distinct lipid constraints on flaviviral output compared with mammalian cells.

In summary, our study establishes host-derived lipids as central regulators of cellular state in sterol-auxotrophic tick cells and reveals an unexpected link between lipid availability and viral output. We show that cholesterol and linoleic acid act as permissive signals that license proliferation, and that restricting lipoprotein availability constrains extracellular TBEV RNA accumulation. These findings support a model in which tick cells integrate environmental lipid cues to transition between non-proliferative and proliferative states, with direct consequences for pathogen production. More broadly, our work positions host lipid homeostasis as a functional axis influencing vector–virus interactions and suggests that metabolic state, rather than receptor availability alone, may limit flavivirus output in this system. Given the availability of pharmacological agents targeting cholesterol trafficking and lipid sensing pathways, these findings provide a conceptual framework for exploring lipid modulation as a strategy to interfere with virus production. Together, this work links nutrient sensing, intracellular lipid handling, and pathogen biology into a unified model with implications for vector biology and infectious disease.

## 4. Material and Methods

### 4.1. Sterol conjugation on BSA

Ten mg of each of the sterols cholesterol (Sigma-Aldrich, C8667), ergosterol (Sigma-Aldrich, E6510), and sitosterol (Sigma-Aldrich, 43623) were dissolved in 1 mL absolute ethanol (VWR Chemicals). The 27-OH-CTL was synthesized as described previously ^57^, and dissolved in CHCl_3_ as a 5mM stock. Bovine serum albumin (BSA) (Sigma-Aldrich, A3059) (400 mg) was dissolved in 8 mL of M1 medium (150 mM NaCl, 5 mM KCl, 1 mM CaCl_2_, 1 mM MgCl_2_, 5 mM Glucose, 20 mM Hepes, pH 7.3). A solution of BSA in M1 medium (8 mL) was mixed with 80 μL of sterol dissolved in ethanol, vortexed for 5 min, and incubated at room temperature for 1 h. The final solution was filter-sterilized (filter 0.22 μm). The solution was mixed with culture medium in a ratio of 1:9. The final concentration of cholesterol in the medium was 10 μg/mL (25.86 μM).

### 4.2. Ligand blot assessment of cholesterol coupling to BSA

Drops (2 μL) of diluted (15× – 50×) serum or cholesterol/BSA solution were loaded on to a nitrocellulose membrane (Cytiva Whatman). Drops were allowed to dry for 10 min. The membrane was washed with phosphate-buffered saline (PBS) + Tween (final concentration 0.05%). The membrane was then incubated with filipin III (50 μg/mL) (Sigma-Aldrich, F9765) for 20 min, washed with PBS + Tween, and bound cholesterol was visualized through filipin fluorescence by a ChemiDoc system (BioRad) with an exposure time of 0.1s. The membrane was then incubated with Amido Black (Sigma-Aldrich) at a final concentration of 0.1% in 10% acetic acid for 15 min and destained with destaining solution (40% methanol, 10% acetic acid). Proteins were visualised in the colorimetry mode of the ChemiDoc (BioRad).

### 4.3. Gas chromatography

Four independent 5mg samples of unfed *I. ricinus* larvae, nymphs, adult females and eggs from a laboratory colony maintained at the Institute of Parasitology, Czech Academy of Sciences, and cultured cell pellets were used for GC-MS analysis. The samples were derivatised with N-trimethylsilylimidazole (TMSI) and saponified as previously described^58^. Briefly, after mixing with 250 µL of 2 M KOH in 90% EtOH (120 min, 60 °C), and centrifugation (4 °C, 10 min, 9,500 × *g*), the samples were extracted with 500 µL of n-hexane and the 50 µL top layer was transported to the derivatization vial. The internal standard 5α-cholestane (total 5 µg per sample) was added to the extract and evaporated to dryness in a nitrogen stream. Derivatization was performed according to the following procedure: 100 µl dimethylformamide and 30 µL TMSI were added to each sample and then heated at 80 °C (30 min). Finally, the TMSI derivatives of cholesterol and the internal standard were extracted from the reaction medium using 100 µL of isooctane.

After mixing and waiting 1 min for the layers to separate, the isooctane layer (upper layer) was transferred to a new vial. One µL was injected into 7890B, a gas chromatograph combined with 1070B mass spectrometer (both Agilent, Santa Clara, CA, USA). Cholesterol was separated using a 30 m VF1-ms (0.25 mm × 0.25 µm thickness) capillary (Agilent, Santa Clara, CA, USA). GC/MS settings were: helium flow rate 1.1 mL/min; injector temperature 280 °C; injection mode splitless; split flow 30 mL/min; splitless time 0.7 min; temperature program 100 °C, 15 °C/min to 320 °C, 3 min hold; transfer line temperature 280 °C; and electron source temperature 230 °C, ionisation energy, 70eV. The mass spectrometer was operated in full scan mode (m/z 40 – 600). Data were acquired and processed using Agilent MassHunter Version 10 software (Agilent, Santa Clara, CA, USA).

### 4.4. Cell culturing and preparation for confocal microscopy

Cells of the *I. ricinus* embryo-derived cell line IRE/CTVM19^17^,sourced from the Tick Cell Biobank, University of Liverpool, were seeded in sealed flat sided 10 mL tubes (Nunc, 156758) in Leibovitz’s L-15 medium (Biosera, LM-L1050) supplemented with 20% FBS (Biosera, FB-1001/100), 10% tryptose phosphate broth (Thermo Fisher Scientific, 18050039), 1% glutamine stable 100× (Biosera, XC-T1755), 1% antibiotic - antimycotic 100× (Biosera, XC-A4110) and incubated at 28 °C. Medium (1/3 volume) was replaced weekly and cells were split equally into two daughter cultures at intervals of at least 14 days. For culturing cells under different lipid supplementation, FBS was replaced with lipoprotein-deficient human serum (LPDS, Sigma Aldrich, S5519) or full human serum (FS, Sigma Aldrich, H4522). All sera were heat-inactivated (56 °C, 45 min) before use. Vero E6 cells^59^ (African green monkey kidney epithelial cells) were grown in Dulbecco’s modified Eagle’s medium (DMEM) supplemented with 10% FBS, 1% antibiotic - antimycotic 100× (Biosera, XC-A4110) and 1% glutamine stable 100× (Biosera, XC-T1755), and cultured at 37 °C under 5% CO_2_. Cells were split 3 times per week.

For filipin and CellBrite Fix 640 co-staining, tick cells were seeded onto coverslips coated with poly-D-Lysine (Sigma Aldrich, P7886) for 1 h at 28 °C. Vero cells were seeded onto coverslips and incubated overnight in 24-well plates at 37 °C under 5% CO_2_. After three washes with PBS, both cell lines were stained with CellBrite Fix 640 (Biotium, 30089A). The dye was initially diluted 1:1000 in DMSO to prepare storage aliquots at-20 °C and further diluted 1:1000 in PBS before staining. Cells were incubated with the staining solution for 45 min, followed by washing 3× with PBS. Cells were subsequently fixed with 3% formaldehyde for 20 min, washed 3× with PBS, quenched with 50 mM glycine for 30 s, stained with 50 µg/mL filipin (Fermentek, FLP-001) for 2 h, washed 3× with PBS and mounted in Fluoromount (Sigma-Aldrich, F4680). The fluorescence signal was recorded with an FV3000 Olympus confocal microscope and images processed by Olympus FV31S-SW software.

For filipin and LD540^60^ (Enamine) or LysoTracker™ Red DND-99 (ThermoFisher Scientific, L7528) co-staining, tick cells were seeded on poly-D-Lysine-coated coverslips for 1 h at 28 °C. After washing 3× with PBS cells were fixed with 3% formaldehyde for 20 min, washed 3× with PBS, quenched with 50 mM glycine for 30 s, stained with 50 µg/mL filipin complex (Fermentek, FLP-001) for 2 h, washed 3× with PBS, stained with LD540 or LysoTracker (1:10,000) for 30 min, washed 3× times with PBS and mounted in Fluoromount (Sigma-Aldrich, F4680). The fluorescence signal was recorded with an FV3000 Olympus confocal microscope and images processed by Olympus FV31S-SW software.

### 4.5. Co-culturing of cells with inhibitors

IRE/CTVM19 cells were seeded in flat-sided tubes and maintained for 14 days without medium exchange, then pre-starved of lipids for 5 days in complete L-15 medium containing LPDS instead of FBS, and then incubated for 7 days in complete L-15 medium containing FS instead of FBS, and supplemented with 50 µM chloroquine diphosphate salt (Sigma-Aldrich, C6628) dissolved in H_2_O, or with 1 µM U18666A (Merck, U3633) dissolved in DMSO. Filipin and CellBrite Fix staining was performed as described above.

Vero E6 cells were seeded overnight onto coverslips in 24-well plates at 37 °C under 5% CO_2_ and then U18666A (Sigma-Aldrich, U3633) was added to a final concentration of 1 µM. After 2 days, cells were washed 3× with PBS, fixed with 3% formaldehyde for 20 min, washed 3× with PBS, quenched with 50 mM glycine for 30 s, stained with 50 µg/mL filipin complex (Fermentek, FLP-001) for 2 h, washed 3× with PBS, stained with LysoTracker (1:10 000) for 30 min, washed 3× with PBS and mounted in Fluoromount (Sigma-Aldrich, F4680). The fluorescence signal was recorded with an FV3000 Olympus confocal microscope and images processed by Olympus FV31S-SW software.

### 4.6. BSA-fatty acid supplementation of tick-cell cultures

IRE/CTVM19 cells were cultured in LPDS-supplemented L-15 medium and supplemented with BSA-conjugated fatty acids. The fatty acid-BSA complexes (Cayman Chemical) used were BSA-linoleate (No. 38649), BSA-palmitate (No. 29558), BSA-arachidonate (No. 34931), BSA-oleate (No. 29557), and BSA-γ-linolenate (No. 39149), together with the corresponding fatty acid-free BSA control (No. 29556). Five-millimolar fatty acid stocks were diluted five-fold in 150 mM NaCl, pH 7.4, to obtain equalised 1 mM stocks. These were further diluted in M1 medium to a concentration of 150 μM. For proliferation assays, cells were seeded at 2.7 × 10^5^ cells per milliliter in LPDS-supplemented medium treated with fatty acid-BSA complexes at a final fatty acid concentration of 15 μM. Control cultures received the corresponding BSA control. Cultures were maintained for 5 days under standard conditions for IRE/CTVM19 incubation, after which cell numbers were determined for 1 mL. For combined lipid supplementation, cells were cultured in LPDS medium containing BSA-conjugated cholesterol, BSA-conjugated linoleate, or both, at final concentrations of lipid ligands approximately 22 μM cholesterol and 15 μM fatty acid.

### 4.7. Sub-cellular localisation of free cholesterol by confocal microscopy

Confocal imaging of IRE/CTVM19 and Vero E6 cells on coverslips was performed using an FV3000 laser scanning microscope (Olympus) equipped with the 100× NA 1.4 oil objective and spectral GaAsP detectors. LD540 (Enamine) and LysoTracker Red were observed with excitation at 561 nm and emission at 570–620 nm, and filipin with excitation at 405 nm and emission at 430–470 nm. Image pixel size was set using the Nyquist sampling criterion. Microscopy images were processed using the Fiji software. To evaluate the distribution of filipin in the plasma membrane and cytoplasm, the background was subtracted from the images and selection of cell outlines was done manually using differential interference contrast images of cells. The plasma membrane was set as a band of 0.8 μm from the cell border. The nucleus was selected using the magic wand (tolerance 100 pixels, 4-connected mode). The cytoplasmic region was set by using the area inside the plasma membrane band excluding the nucleus. Mean fluorescence intensity values were measured for the plasma membrane and cytoplasm regions and corrected for autofluorescence values in the respective regions. For colocalization analysis, cells were segmented using Cellpose v4.0.6 (SAM-Segment Anything Model)^61^, thresholding, and manual adjustments. Then, using Python code for calculating Pearson’s correlation coefficient, modelled after the Coloc2 plugin in Fiji, the colocalization of features was measured across the two fluorescence channels for each cell, excluding the mask of the nuclei. Cells that were partially cut off by the edges of the image were excluded. Statistical analysis of the data was done using GraphPad Prism. Normality of the data distribution was tested and verified using D’Agostino-Pearson omnibus normality test. Multiple comparisons were done using Brown-Forsythe and Welch ANOVA tests with Dunnett T3 correction. A *P* value of less than 0.05 was considered statistically significant.

### 4.8. Preparation of LIMP2 - mCherry fusion construct

A fusion construct connecting mCherry fluorescent protein to the C terminus of LIMP-2 via a short flexible linker (GSA GSA AGS GEF)^62^ was designed using the NEBuilder® online assembly tool (NEB). The gene encoding LIMP2 from the hard tick *I. ricinus* without a stop codon was amplified using LIMP2-C-F and R primers (Supplementary Table S2). The gene encoding mCherry without its start codon was amplified using mCher-C-LIMP-F and R primers. The LIMP2-C-R and mCher-C-LIMP-F primers included the sequence for the flexible linker. Amplified LIMP2 and mCherry fragments were assembled into the pUC19 cloning vector (amplified using pUC19F and R primers) via the NEBuilder^®^ HiFi DNA Assembly Master Mix (NEB) according to the manufacturer’s instructions and the sequence was verified by Sanger sequencing. The resulting fusion gene was reamplified with Limp-C-mCher-DES-F and R primers, and inserted into the pMT/V5-hisA expression vector (Thermo Fisher Scientific) using XhoI and AgeI restriction enzymes and T4 DNA ligase. Sequence-verified expression vector carrying LIMP2 was purified by a NucleoBond Xtra maxi kit (Macherey Nagel), and stored at-20 °C prior to transfection into S2 cells.

### 4.9. LIMP2 production in S2 cells and lysosome co-localization

*Drosophila* S2 cells^63^ were grown in reconstituted Schneider′s *Drosophila* powdered medium (Serva, cat. No. 47521.04) supplemented with 8% FBS (Biosera) and 1% of 100× penicillin/streptomycin solution (Biosera) (S-complete) in sealed T25 cell culture flasks (TPP) at 27 °C. For transfection experiments, the Escort™ IV transfection reagent (Merck) was used according to the manufacturer’s instructions. Aliquots of 1x 10^6^ cells were seeded into 12 well plates (TPP) in 1 mL of S-complete and allowed to grow for 24 h. Prior to the transfection, S2 cells were washed with 1mL of Schneider′s Insect Medium without FBS and antibiotics and 400 µl of the same medium was added. Next 2 µg of plasmid DNA was added to 50 μL of Schneider′s Insect Medium, mixed with Escort medium (3µL of Escort transfection reagent and 47 μL Schneider′s Insect Medium), and incubated at room temperature for 45 min. The resulting DNA complexes were added dropwise to the S2 cells and transfected for 20 h at 27 °C. The transfection medium was then replaced with 1 ml of fresh S-complete and protein production was induced with 500 µM CuSO_4_ (final concentration). The cells were incubated at 27 °C for three days prior to the next step. Transfected *Drosophila* S2 cells (1 mL; 1 × 10⁶ cells) were seeded onto poly-D-lysine–coated coverslips (Sigma-Aldrich, P7886) in a 24-well plate and allowed to attach for 1.5 h at 28 °C. LysoTracker DND-26 (Thermo Fisher Scientific, L7526) at 75 nM was then added, and cells were incubated with the staining solution for 30 min. Hoechst 33342 (Thermo Fisher, 62249) was diluted to a 2 µM solution in PBS and subsequently added to counterstain cell nuclei. Cells were washed twice with PBS and mounted in Fluoromount (Sigma-Aldrich, F4680). Fluorescence signals were recorded using an Olympus FV3000 confocal microscope, and images were processed with Olympus FV31S-SW software.

### 4.10. RNA isolation and RT-qPCR or RNA-Seq data processing

Tick cells were harvested and total RNA was isolated using Nucleospin RNA (Macherey-Nagel). Single-stranded cDNA was reverse-transcribed from 0.5 µg of total RNA using the Transcriptor High-Fidelity cDNA Synthesis Kit (Roche Diagnostics, Germany). For subsequent RT-qPCRs, cDNA was diluted 20 times in nuclease-free water. Samples were analysed using a LightCycler 480 (Roche) and Fast Start Universal SYBR Green Master Kit (Roche). Each primer pair (for the list of qPCR primers, see Supplementary Table S2) was inspected for its specificity using melting curve analysis. Relative expressions of tick mRNA transcripts were calculated using the ΔΔCt method, normalised to *elongation factor 1* of *I. ricinus* (*ef-1α*; GeneBank ID: GU074828), using previously verified primers ^64^. The RNA was also sent for Illumina sequencing at Novogene. After passing the quality check, sequencing was performed on an Illumina NovaSeq platform using a paired-end 150 bp (PE150) configuration. Approximately 50 million raw reads were generated per library. Across all samples, raw read counts ranged from ∼49.9 to 67.1 million reads per library. Base quality metrics demonstrated high sequencing quality, with Q20 scores ranging from 96.33% to 97.18% and Q30 scores from 90.66% to 92.35%. The RNA-Seq data quality was monitored using FastQC (version 0.11.9; http://www.bioinformatics.bbsrc.ac.uk/projects/fastqc/), and reads were trimmed and filtered using fastp (version 0.23.2) ^65^ to remove remaining adapter sequences as well as bases with a quality lower than 30 from both the 5’and 3’ ends. Reads with more than two N bases or a length of less than 15 bases after the trimming step were filtered from the dataset.

### 4.11. *De novo* transcriptome assembly and evaluation

For the *de novo* transcriptome assembly, all 15 pairs of RNA-Seq datasets were given as input for Trinity (version 2.14.0) ^66^ using standard parameters. The resulting assembled contigs were clustered to remove redundant contigs by applying Linclust from MMSeqs2 (version 13.45111)^67^. To evaluate the quality of the assembled non-redundant transcriptome, two independent measures were used: the back-mapping rate and the BUSCO completeness score.

The individual RNA-Seq dataset pairs were mapped to the assembled transcriptome to calculate the back-mapping rate using HISAT2 (version 2.2.1) ^68^. To estimate the completeness of the assembled transcriptome we used BUSCO (version 5.3.2) ^69^ based on the arachnida_odb10 database.

### 4.12. Transcriptome annotation and quantification

To ascribe potential functions to the assembled transcripts, two different approaches were used. The first approach was based on the tool MetaEuk (version 5.34c21f2)^70^, to identify potential encoded protein sequences and subsequently compare these predicted protein sequences with the NCBI invertebrate reference sequence database (RefSeq release 213), to identify their potential function using MMSeqs2. We considered the protein prediction of an assembled transcript to be a homolog of a RefSeq protein if the corresponding hit had an e-Value of less than 1e-20, if the two proteins had at least 80% sequence identity and if the predicted protein had at least 70% of the respective RefSeq length of protein. In the second approach, we performed homology comparisons of the assembled transcripts with the mRNA sequence dataset of *I. scapularis* using blastn (version 2.12.0+)^71^. We considered an assembled transcript to be a homolog of an *I. scapularis* mRNA if the hit had an e-Value of less than 1e-10, if the RNAs had at least 90% sequence identity, and if the assembled transcript had at least 90% of the respective *I. scapularis* mRNA sequence length. For transcript quantification, we used the mapping results produced by HISAT2 from the back-mapping evaluation step. For read counting, FeatureCounts (version 2.0.1)^72^ with the parameters - p for fragment counting,-M for multimapping, and-s 0 for strand-unspecific counting was used. Based on these raw count values, FPKM and TPM values were calculated. Differentially-expressed transcripts were identified by filtering ≥ 2.5-fold changes in both full serum or BSA-Chol supplementations in a given time-point group.

### 4.13. Virus infection

Tick-borne encephalitis virus (TBEV) strain Neudoerfl was used throughout the study. This strain was originally isolated from *I. ricinus* ticks in Austria in 1971 ^73^ and was kindly provided by Professor F. X. Heinz (Institute of Virology, Medical University of Vienna, Austria). Prior to use, the virus had undergone five passages by intracranial inoculation of suckling mice.

IRE/CTVM19 tick cells were pre-starved of lipids for 8 days in complete L-15 medium containing LPDS instead of FBS in flat-sided culture tubes, and then seeded in 0.5 × 10^6^ cells per well of 24-well plate and incubated for 1 day in the medium with LPDS, followed by exchange of medium containing LPDS or FS (6 wells per diet)before infection with TBEV at an MOI of 2. Virus adsorption was carried out for 6 h at 28 °C in 0.5% CO_2_. Following adsorption, the cells were washed five times with medium and resuspended in fresh medium containing the corresponding lipid supplement (FS or LPDS). Cells were then incubated for 3 days at 28 °C in 0.5% CO_2_ in a total volume of 1 mL. At the end of the incubation period, culture supernatants were collected, stored at-80°C, and used for RNA isolation. Cell pellets were resuspended in MagMAX mirVana lysis buffer (Thermo Fisher Scientific) and stored at-80 °C until further processing.

### 4.14. Isolation of total RNA and estimation of TBEV viral genome copies

Total RNA was extracted from harvested cell pellets from each well and from 200 µL of culture supernatant using the MagMAX™ mirVana™ Total RNA Isolation Kit (Thermo Fisher Scientific) on a KingFisher™ Apex system (Thermo Fisher Scientific), according to the manufacturer’s instructions. For cDNA synthesis, 5 µL of isolated RNA was reverse-transcribed using the High-Capacity RNA-to-cDNA Kit (Thermo Fisher Scientific) following the manufacturer’s protocol. TBEV genome copies were quantified from 5 µL of the resulting cDNA as described previously ^74,75^, using SsoAdvanced™ Universal SYBR® Green Supermix (Bio-Rad) according to the manufacturer’s instructions. A synthetic single-stranded DNA standard was used to generate a calibration curve for the quantification of viral genome copy numbers in the samples.

For host gene expression analysis, 5 µL of synthesised cDNA was subjected to SYBR Green-based real-time PCR using SsoAdvanced™ Universal SYBR® Green Supermix (Bio-Rad). Transcript for *ef-1α* served as the endogenous reference tick transcript used for normalisation. Relative TBEV RNA levels were determined using the comparative Ct method and expressed as fold change relative to the control condition. Briefly, ΔCt values were calculated by subtracting the Ct value of *ef-1α* from that of the target gene. ΔΔCt values were then determined as ΔCt of experimental samples (LPDS) minus ΔCt of the control samples (FS). Fold changes were calculated as 2^-ΔΔCt, as previously described ^76^. Statistical analysis of viral expression was performed using Mann-Whitney test in GraphPad Prism 10.6.1. A *P* value of less than 0.05 was considered statistically significant.

## 5. Acknowledgement

The authors thank Eva Výletová and Marika Davídková from the Laboratory of Arbovirology for excellent technical support.

## 6. Data availability

The cellular images from confocal microscopy are available at BioImage Archive. The accession number is S-BIAD3266. The RNA-seq data has been deposited at the NCBI server, with the Geo Accession ID: GSE317082 (will be available upon publication). The Mass Spectrometry data are available at: https://doi.org/10.6084/m9.figshare.31241068.

